# Mycn regulates vascular development through PI3K signaling pathway in zebrafish

**DOI:** 10.1101/2024.08.19.608563

**Authors:** Guo-Qin Zhao, Tao Cheng, Peng-Yun Wang, Jing Mo, Feng Yu, Yang Dong, Yun-Fei Li, Yu Feng, Peng-Fei Xu, Li-Ping Shu

## Abstract

Mycn, a MYC gene family member, is implicated in both carcinogenesis through amplification and Feingold syndrome through its deficiency. Previous studies have indicated that increased Mycn expression enhances vascularization in human neuroblastomas, yet its precise role in vascular development remains elusive. In this study, we utilized single-cell RNA-seq and live imaging analyses to confirm that *mycn* is expressed during zebrafish vasculogenesis. We investigated vascular development in zebrafish using a genetically engineered *mycn* mutation. Our findings reveal that *mycn*-deficient zebrafish exhibit reduced intersegmental vessels and malformed subintestinal vessels, primarily due to decreased cell proliferation in vascular cells. Importantly, we discovered that activation of PI3K signaling significantly ameliorates these vascular abnormalities.

## Introduction

Mycn, a highly conserved transcription factor in vertebrates, belongs to the MYC proto-oncogene family^1–2^. It plays a pivotal role in early embryonic development, with predominant expression during the embryonic stage and involvement in regulating key cellular processes such as proliferation, differentiation, apoptosis, protein synthesis, and metabolism^3^. In mice, the genetic deletion of Mycn results in embryonic lethality by day E11.5, impacting the development of critical organ systems including the nervous system, heart, lungs, gut, and cardiovascular system^4^. Furthermore, Mycn is linked to the development of several childhood diseases, predominantly tumors^5^. Studies on the Mycn gene have largely focused on neuroblastoma, where its amplification or overexpression correlates strongly with the tumor’s prognosis^6–9^. Additionally, increased vascularization, influenced by Mycn, plays a crucial role in the progression of neuroblastoma^10–11^.

The vascular system is a crucial component in all vertebrates, responsible for delivering oxygen and essential nutrients to every tissue and organ^12^. Many congenital disorders are linked to defects in vascular formation and cardiovascular development^13^. Angiogenesis, the physiological process through which new blood vessels form from pre-existing ones^14^, is central to both normal organ growth and tumor progression^15–16^. This process is tightly regulated by a balance of proangiogenic and antiangiogenic genes. While essential for normal development, tissue repair, and organ regeneration, angiogenesis can also lead to pathological conditions, including the advancement of cancer. Tumor angiogenesis is pivotal in controlling tumor growth and metastasis, underscoring its role in cancer progression^17–18^.

Zebrafish present significant advantages for investigating vascular development due to the conserved developmental processes and regulatory molecular mechanisms with other vertebrates, including humans^19–20^. Additionally, the transparent embryos, rapid development and availability of genetic manipulation make zebrafish act as an ideal research model to study angiogenesis *in vivo*^21–22^.

In this study, we explored the role of Mycn in regulating angiogenesis in zebrafish through Mycn loss-of-function. We found that Mycn deficiency leads to the reduction of intersegmental vessels and malformation of subintestinal vessels in zebrafish embryos during hatching stage. Notably, the angiogenesis defects in *mycn* mutants result from impaired cell proliferation rather than apoptosis. By conducting single-cell RNA-seq at 72 hpf and bulk RNA-seq analyses at 48 hpf on wild-type and *mycn* mutant embryos, we identified a potential involvement of the PI3K/mTOR signaling pathway in the angiogenesis defects resulting from Mycn deficiency. Supplement of L-Leucine, known to enhance mTOR signaling activity, partially rescued the angiogenesis defects. However, treatment with 740Y-P, an activator of the PI3Ksignaling, significantly improved both intersegmental and subintestinal vessel abnormalities in *mycn* mutants.

## Results

### *mycn* is expressed in the vascular endothelial cells of zebrafish embryos

By analyzing the single-cell RNA-seq datasets of wild-type embryos during early development from our previous studies and other studies^23–24^, we observed that Mycn is extensively expressed in zebrafish embryonic blood vessel cells (Figs. 1A and S1F-H). To validate these observations, We performed live imaging on Ki(*mycn*:EGFP);Tg(*kdrl*:mCherry) embryos, which were generated by crossing the Ki(*mycn*:EGFP) and Tg(*kdrl*:mCherry) zebrafish reporter lines. No *mycn*^+^*kdrl*^+^ cells were observed at 24 hours post fertilization (hpf) (Fig. S1A). However, from 30 to 72 hpf, we detected *mycn*^+^*kdrl*^+^ cells in the dorsal aorta, caudal artery, and caudal vein plexus (Figs. S1B-S1E). Remarkably, at 30 hpf, the *mycn*^+^*kdrl*^+^ cells were predominantly located on the ventral wall of the dorsal aorta and peaked in number by 36 hpf. Subsequently, these cells mainly located in the caudal dorsal artery and caudal venous plexus. Interestingly, we noted that these *mycn*^+^*kdrl*^+^ cells exhibited either a flattened or spherical shape (Figs. 1B), and some engaged in pseudopodial protrusion to facilitate migration or underwent cell division (Figs. 1C, Movies 1-3). These observations suggest that Mycn may function as a regulator of vascular development.

**Figure 1.**
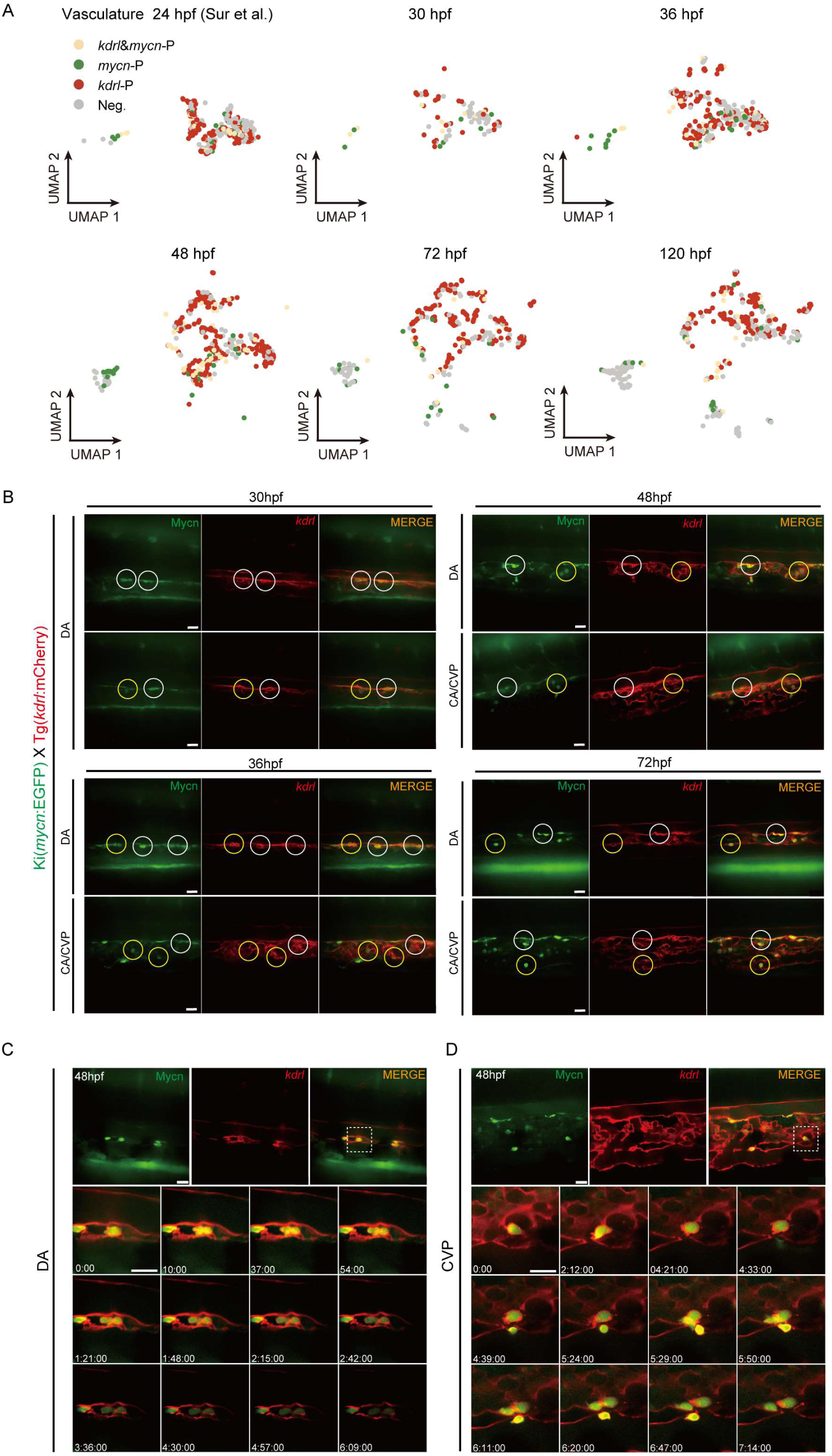
The cellular behavior of *mycn*^+^*kdrl*^+^ cells. (A) Visualizing *kdrl*-positive and *mycn*-positive cells on the UMAP of zebrafish vasculature at 24, 30, 36, 48, 72 and 120 hpf. (B) Live images showing the morphology of *mycn*^+^*kdrl*^+^ cells at 30, 36, 48 and 72 hpf. The *mycn*^+^*kdrl*^+^ cells in white circle display flattened morphology, while *mycn*^+^*kdrl*^+^ cells in yellow circle display spherical shapes. (C) Time-series of the DA from a Ki(*mycn*:EGFP);Tg(*kdrl*:mCherry) embryo from 48 to 55 hpf. The *mycn*^+^*kdrl*^+^ cell that underwent cell division was captured in the DA. (D) Time-series of the CVP region from a Ki(*mycn*:EGFP);Tg(*kdrl*:mCherry) embryo from 48 to 56 hpf. The *mycn*^+^*kdrl*^+^ cell displayed pseudopodial movement and occasional cell division in the CVP. Scale bars: 20 μm.

### The development of ISV and SIV was impaired in *mycn* mutants

By analyzing single-cell RNA-seq data of *mycn* mutant embryos from our previous studies^23^, we observed a reduction in blood vessel cells in *mycn* mutant compared to wild-type embryos at 72 hpf (Fig. 2A and S2). To validate this observation, we perform WISH for *kdrl*, a vascular endothelial cell marker^25^, in wild-type and *mycn* mutant embryos at 48 and 72 hpf. The results showed a dramatic decrease in *kdrl* expression in *mycn* mutants, indicating a significant vascular development defect due to the *mycn* mutation (Fig. 2B).

**Figure 2.**
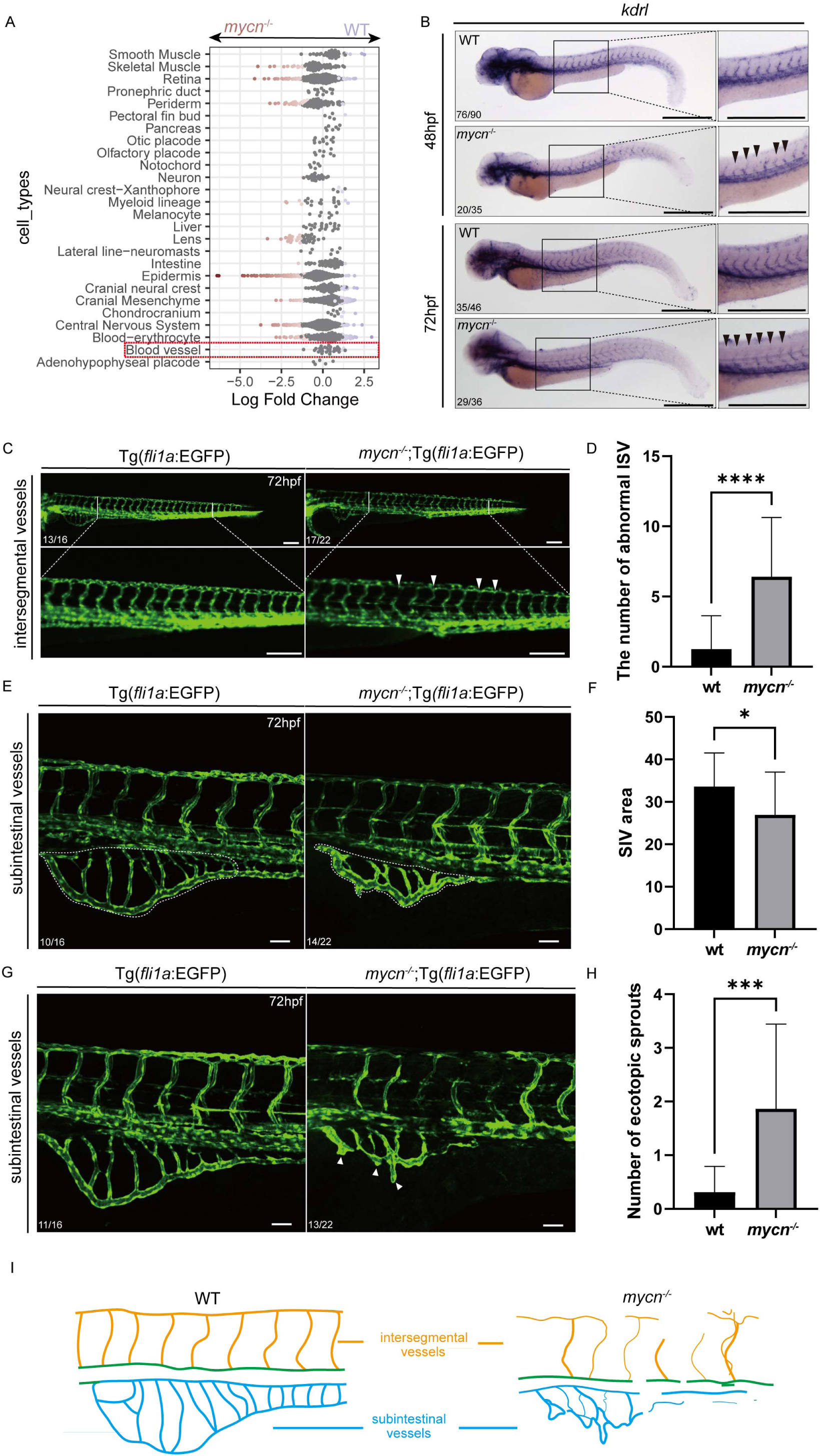
Mycn deficiency leads to abnormalities in the formation of intersegmental vessels and subintestinal vessels. (A) Beeswarm plot showing the differential abundance by cell types between wild-type and *mycn*-mutant embryos. (B) WISH images showing *kdrl* expression in wild-type and *mycn*-mutant embryos at 48 hpf and 72 hpf. The panel on the right is a magnified image of the boxed area on the left. The black arrow indicates the intact ISVs. (C) Fluorescence images showing the ISV morphology of Tg(*fli1a*:EGFP) and *mycn*^-/-^; Tg(*fli1a*:EGFP) zebrafish embryos at 72 hpf. The panel on the bottom is a magnified image of the boxed area at the top. White arrowheads indicate abnormal ISVs. (D) Quantification of the number of abnormal ISVs in (C). (E) Fluorescence images showing the morphology of the whole SIVs in Tg(*fli1a*:EGFP) and *mycn*^-/-^;Tg(*fli1a*:EGFP) embryos at 72 hpf. The white dashed line denotes the range of whole SIVs. (F) Quantification of the area of ISVs in (E). (G) Representative fluorescence images showing the SIV morphology of Tg(*fli1a*:EGFP) and *mycn*^-/-^;Tg(*fli1a*:EGFP) embryos at 72 hpf. White arrowheads indicate the ectopic sprouts. (H) Quantification of the number of ectopic sprouts in (G). (I) Schematic diagram showing the morphology of ISVs and SIVs in Tg(*fli1a*:EGFP) and *mycn*^-/-^;Tg(*fli1a*:EGFP) embryos at 72 hpf. Statistical differences between two samples were evaluated by Student’s t-test (D, F and H). *indicates P-value < 0.05; ***indicates P-value < 0.001; ****indicates P-value < 0.0001. Each experiment was performed for at least 3 independent replicates (technical replicates). Scale bars: 500 μm (B), 200 μm (C) and 50 μm (E and G).

To further explore these vascular developmental defects in *mycn* mutants, we focused on two types of vessels: intersegmental vessels (ISVs) and subintestinal vessels (SIVs). We crossed the *mycn* mutants with Tg(*fli1a*:EGFP) lines, and further obtained homozygous *mycn* mutants with Tg(*fli1a*:EGFP) transgenic background. Subsequent imaging revealed that ISVs, which typically develop from the dorsal aorta around 22 hpf and navigate between somites to form the dorsal longitudinal anastomotic vessel^26^, exhibited significant developmental defects in *mycn* mutants by 30 hpf, both in fluorescent intensity and morphology (Figs. 2C-2D). SIVs, originating from the endothelial cells of the posterior cardinal vein and forming vascular plexuses around the intestine from 48 hpf^27^, were also affected. Live imaging of Tg(*fli1a*:EGFP) and *mycn*^-/-^;Tg(*fli1a*:EGFP) embryos during SIV development (Movies S4-S5) showed a significant reduction in the area of SIVs and a large number of ectopic sprouts in *mycn* mutants (Figs 2E-2H). These results strongly suggest that *mycn* deficiency leads to significant abnormalities in the formation of both intersegmental and subintestinal vessels (Fig. 2I).

### Cell proliferation is impaired in the blood vessel of *mycn* mutants

To investigate the cellular mechanisms underlying the angiogenesis defects in *mycn* mutants, we isolated blood vessel cells from single-cell RNA-seq datasets^23^ of wild-type and *mycn* mutant embryos at 72 hpf, and performed re-clustering analyses. We obtained 9 subclusters, annotating them as epidermal (*cldni*, *pfn1*), endothelial progenitor (*etv2*, *flt4*), arterial endothelial (*flt1*, *aqp1a.1*), endocardial (*f8*, *fn1a*), venous endothelial (*dab2*, *lyve1b*), proliferating (*smc2*, *mki67*), lymphatic endothelial (*prox1b*, *cdh6*), mesenchyme (*col6a3*, *fap*), neural (*elavl3*, *scrt2*) separately^28–30^ (Figs 3A and S3A). Interestingly, a comparative analysis revealed a decrease in endothelial progenitor cells and proliferating cells (expressing high levels of proliferation-related genes) in *mycn* mutants compared to wild-type embryos (Figs 3B and S3B). These findings suggest that cell proliferation may be impaired in *mycn* mutants. To further validate this hypothesis, we conducted EdU and TUNEL assays to assess cell proliferation and apoptosis, respectively, in Tg(*fli1a*:EGFP) and *mycn*^-/-^;Tg(*fli1a*:EGFP) embryos, particularly focusing on the trunk region. Imaging and subsequent quantification of proliferative and apoptotic cells revealed a significant reduction in cell proliferation in *mycn* mutants (Fig. 3C-3H), while apoptosis levels did not show significant differences between *mycn* mutants and wild-type embryos (Fig. S4A-S4F).

**Figure 3.**
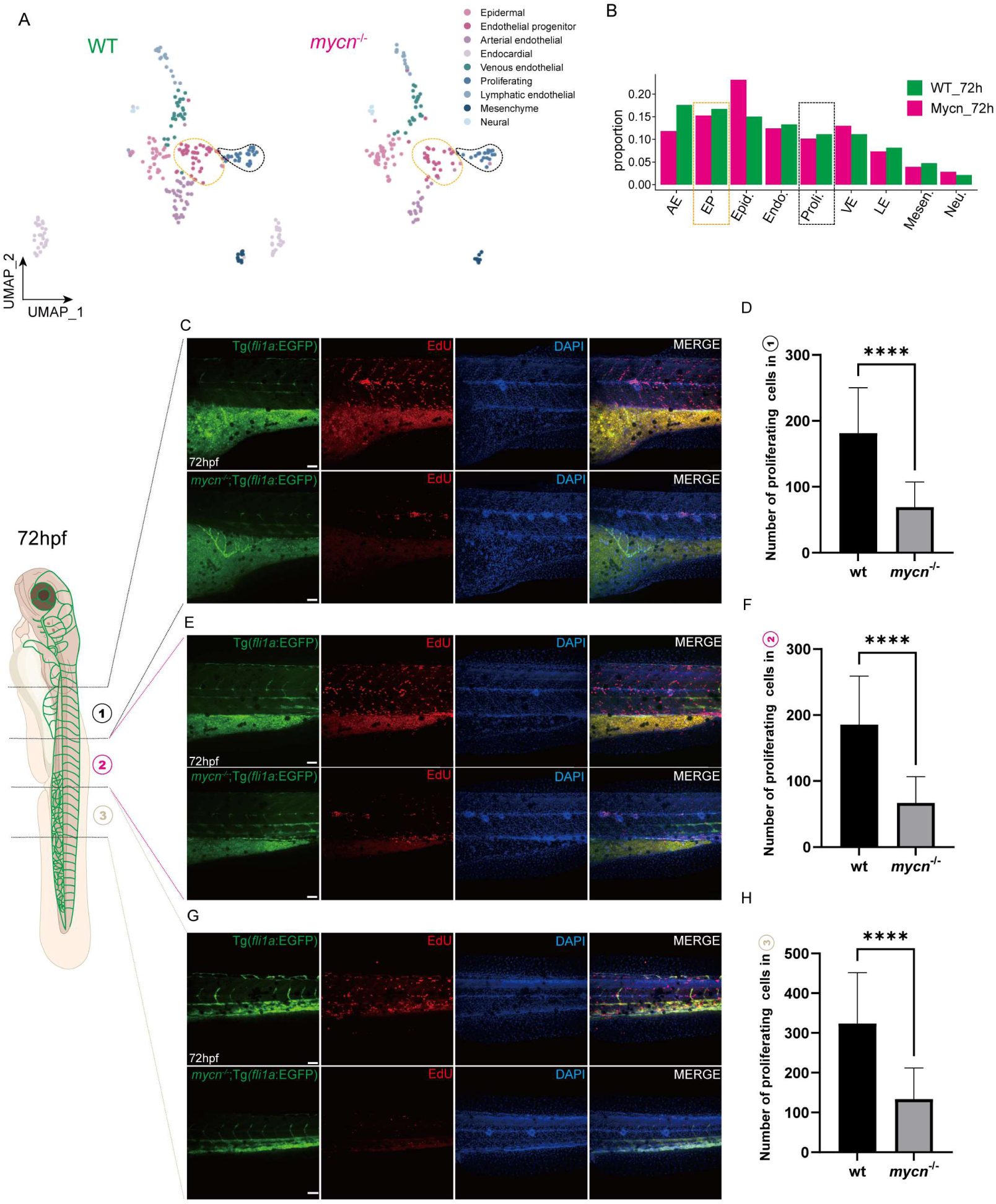
Single-cell RNA-seq analysis and EdU assays showed that proliferation signals were significantly reduced in vessel regions in the *mycn*-mutant embryos. (A) UMAP plot showing the subclusters of blood vessel cells selected from Figure.S2(A). A total of 9 clusters were identified by unsupervised clustering. (B) Bar plot shows the proportions of each cell type in *mycn*-mutant (rose) or wild-type embryos (green). Cell proliferation in blood vessels was detected by an EdU assay (red signal) in Tg(*fli1a*:EGFP) and *mycn*^-/-^;Tg(*fli1a*:EGFP) embryos. The vasculature was divided into three parts: the first part contained SIVs(C), the second region included ISVs (E), and the third part contained CVP (G). The statistical analyses of the proliferating cells in each region are shown on the right (D, F and H). Statistical differences between two samples were evaluated by Student’s t-test (D, F and H). NS indicates P-value >=0.05; *indicates P-value < 0.05; *indicates P-value < 0.05; **indicates P-value < 0.01; ***indicates P-value < 0.001; ****indicates P-value < 0.0001. Each experiment was performed for at least 3 independent replicates (technical replicates). Scale bars: 50 μm.

Taken together, these results indicate that impaired proliferation, rather than apoptosis, contributes to the angiogenesis defects observed in the *mycn* mutants.

### Activation of PI3K signalling rescues the angiogenesis defects in *mycn* mutants

Our previous research demonstrated that increasing mTOR signaling, either through L-Leu treatment or *rheb* mRNA injection, could ameliorate intestinal defects caused by Mycn deficiency in zebrafish^23^. To assess whether mTOR signaling activation could similarly rectify angiogenesis defects in *mycn* mutants, we injected L-Leu into the yolk sac of *mycn* mutant embryos at 30 hpf and examined vascular development at 72 hpf. Although L-Leu slightly improved the angiogenesis defects, this suggested that other signaling pathways might play a more pivotal role in vascular development downstream of Mycn. To identify the potential signalling pathways, we selected blood vessel cells from the single-cell RNA-seq dataset, and performed deferential expression analysis between wild-type and *mycn* mutant embryos at 72 hpf. The down-regulated genes in *mycn* mutants were used for GO enrichments analysis. We found that translation related GO terms were significantly enriched, such as: “cytoplasmic ribosomal proteins”, “ribonucleoprotein complex biogenesis”, “ribosomal small subunit biogenesis” and “translation elongation” (Fig. 4A). These findings align with our earlier discoveries that impaired protein translation contributes to intestinal defects in *mycn* mutants. Remarkably, we found many pik3 genes were dramatically down-regulated in *mycn* mutants (Fig. 4B), including *pik3c3*, whose mutation resulted in intestinal defects in zebrafish similar to that of the *mycn* mutant^31^. Furthermore, previous studies also have reported that the PI3K/mTOR signalling pathway is associated with angiogenesis^32^. To investigate whether PI3K signalling is involved in impaired angiogenesis in *mycn* mutants, we treated the mutants with 740Y-P, which is an activator of PI3K signalling, and then assessed the vascular development in the mutants. We found that the number of abnormal intersegmental vessels was largely decreased in *mycn* mutant embryos after 740Y-P treatment (Figs. 4C-4H), and the subintestinal vessel area and the number of ectopic sprouts were notably rescued.

**Figure 4.**
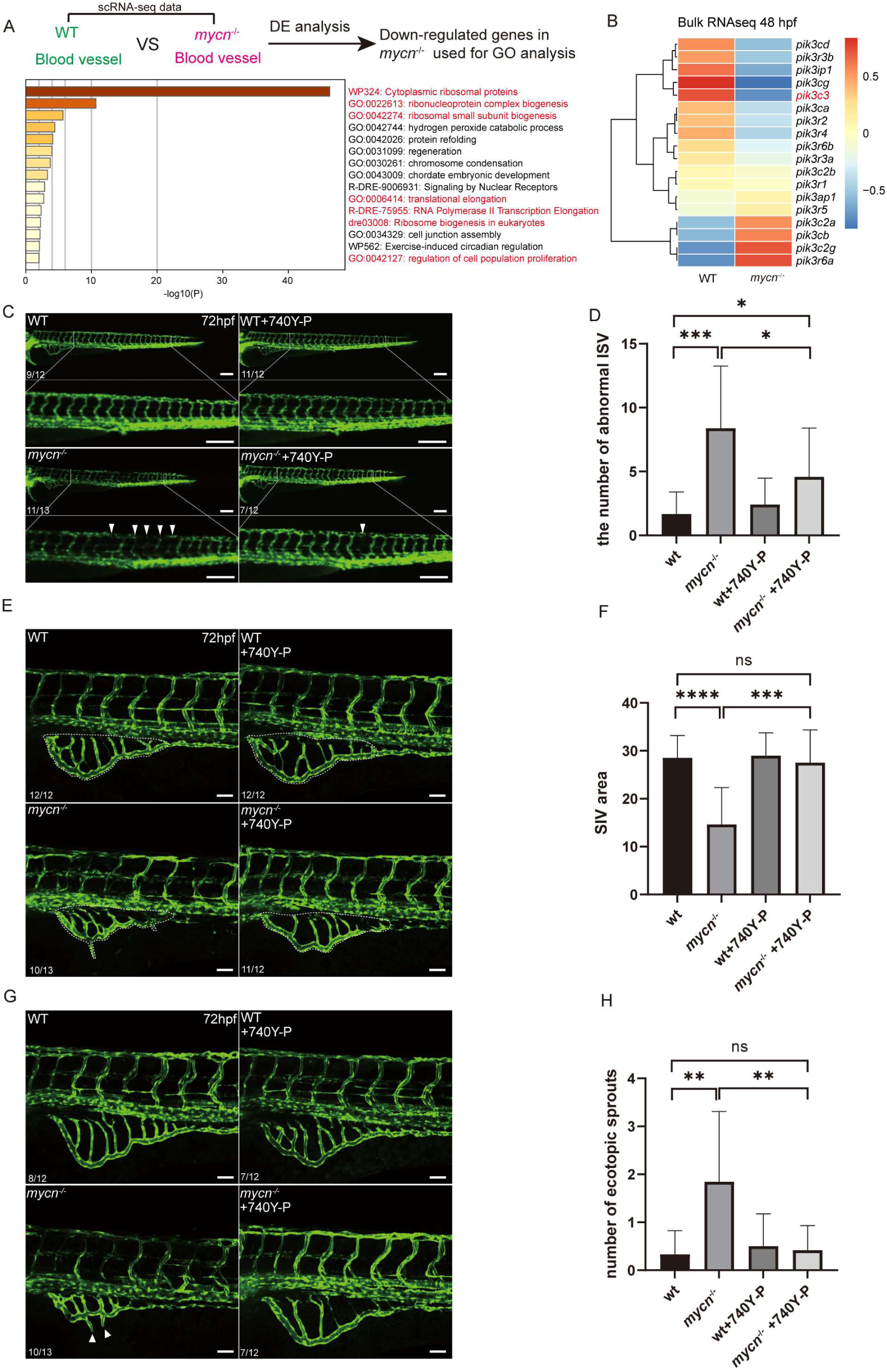
740Y-P treatment dramatically reversed the intersegmental vessel and subintestinal vessel defects. (A) Enriched terms using the down-regulated genes in the *mycn*^-/-^ blood vessel cells. (B) Heatmap showing the expression of PI3K-related genes in WT and *mycn*^-/-^ embryos at 48 hpf. Color indicates Z-score, and each cell represents the mean Z-score of three replicates. (C) Fluorescence images showing the ISVs morphology of 72 hpf Tg(*fli1a*:EGFP) and *mycn*^-/-^;Tg(*fli1a*:EGFP) embryos treated with 740Y-P from 10 to 72 hpf. The panel on the bottom is a magnified image of the boxed area at the top. White arrowheads indicate abnormal ISVs. (D) Quantification of the number of abnormal ISVs in (C). (E) Representative images showing the SIVs morphology of Tg(*fli1a*:EGFP) and *mycn*^-/-^;Tg(*fli1a*:EGFP) zebrafish embryos at 72 hpf after 740Y-P treatment. White dashed line denotes the range of whole SIVs. (F) Quantification of the area of ISVs in (E). (G) Fluorescence images showing the SIVs morphology of Tg(*fli1a*:EGFP) and *mycn*^-/-^;Tg(*fli1a*:EGFP) embryos at 72 hpf with 740Y-P treatment. White arrowheads indicate the ectopic sprouts. (H) Quantification of the number of ectopic sprouts in (G). Statistical differences between two samples were evaluated by Student’s t-test (D, F and H). NS indicates P-value >=0.05; *indicates P-value < 0.05; *indicates P-value < 0.05; **indicates P-value < 0.01; ***indicates P-value < 0.001; ****indicates P-value < 0.0001. Scale bars: 200 μm (C), 50 μm (E and G).

These findings collectively underscore the critical role of PI3K signaling in the formation of intersegmental and subintestinal vessels downstream of Mycn.

### The cerebral vascular development deficiency cannot be rescued by activating mTOR or PI3K signalling pathway

Angiogenesis is a crucial process during brain development in vertebrate embryos, where the proper formation and function of cerebral blood vessels are essential for central nervous system (CNS) development^33^. In *mycn* mutants, we observed abnormalities in the dorsal longitudinal vein (DLV) and the mesencephalic vein (MsV) compared to wild-type embryos: both the length of the DLV and the bifurcation angle of the MsV were significantly reduced at 72 hours post-fertilization (hpf) (Figs. 5A-5B). To explore potential rescue mechanisms, we activated mTOR and PI3K signaling pathways in these mutants by administering L-Leucine (Leu) and 740Y-P, respectively. However, subsequent analyses of cerebral vascular structures revealed that activation of neither mTOR nor PI3K pathways could rescue the cerebral vascular development deficiencies in *mycn* mutants (Figs. 5C-5F). These findings indicate that, unlike in the trunk region, cerebral blood vessel development regulation by Mycn does not substantially rely on the mTOR or PI3K signaling pathways. Therefore, the mechanisms through which Mycn regulates cerebral blood vessel development require further investigation.

**Figure 5.**
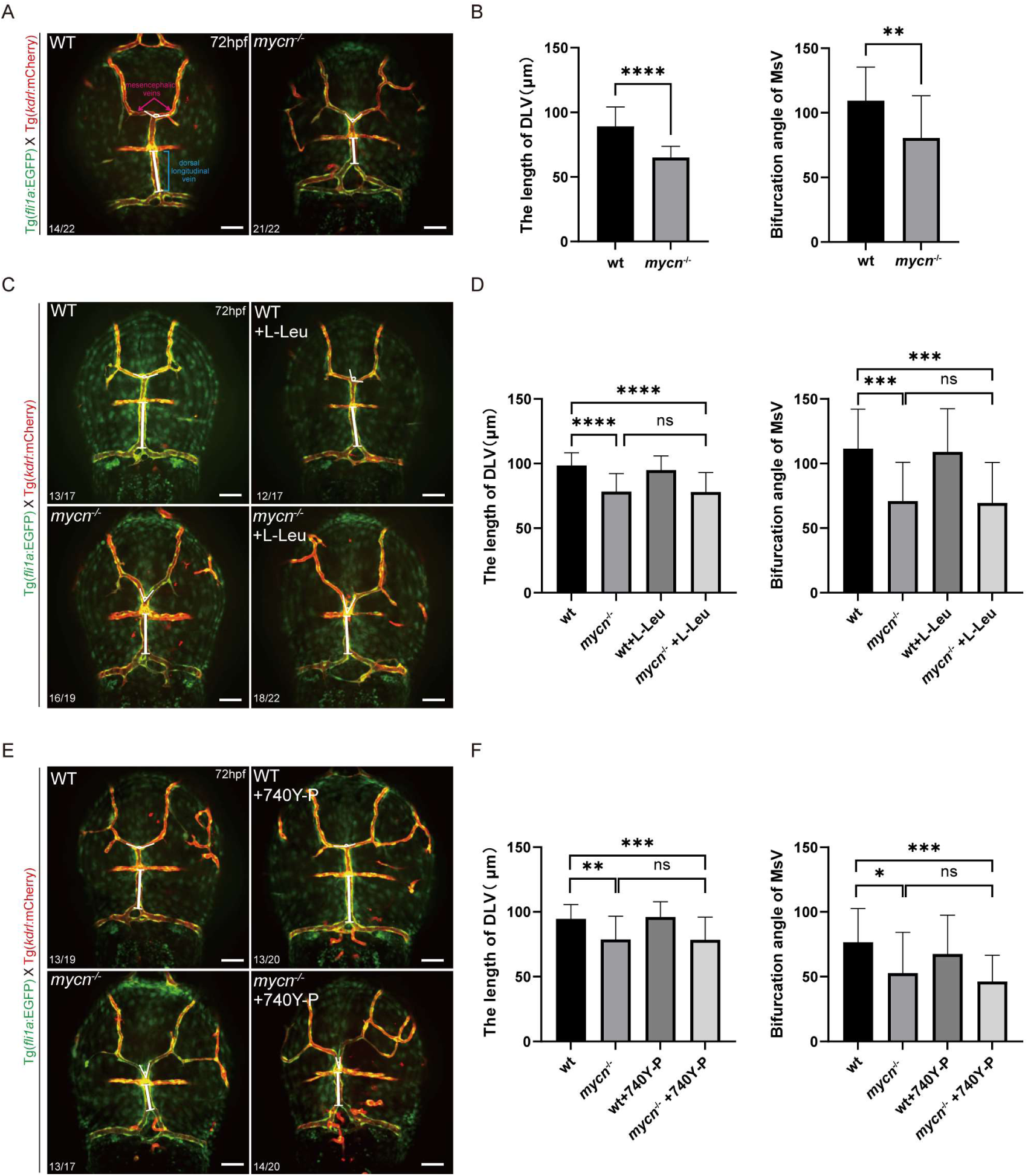
Mycn deficiency caused defects in dorsal longitudinal vein and mesencephalic vein development in the brain. (A) Fluorescence images showing the DLV and MsV morphology of Tg(*fli1a*:EGFP;*kdrl*:mCherry) and *mycn*^-/-^;Tg(*fli1a*:EGFP;*kdrl*:mCherry) zebrafish embryos at 72 hpf. The blue line indicates the DLV, and the rose arrowheads indicate the MsV. (B) The statistical results of the length of DLV and the bifurcation angle of MsV in (A). (C) Fluorescence images showing the DLV and MsV morphology of 72-hpf Tg(*fli1a*:EGFP;*kdrl*:mCherry) and *mycn*^-/-^;Tg(*fli1a*:EGFP;*kdrl*:mCherry) embryos that were injected with L-Leu at 30 hpf. (D) Quantification of the length of DLV and the bifurcation angle of MsV in (C). (E) Fluorescence images showing the DLV and MsV morphology of 72-hpf Tg(*fli1a*:EGFP;*kdrl*:mCherry) and *mycn*^-/-^;Tg(*fli1a*:EGFP;*kdrl*:mCherry) zebrafish embryos treated with 740Y-P from 10 to 72 hpf. (F) Quantification of the length of DLV and the bifurcation angle of MsV in (E). Statistical differences between two samples were evaluated by Student’s t-test (B, D and F). NS indicates P-value >=0.05; *indicates P-value < 0.05; *indicates P-value < 0.05; **indicates P-value < 0.01; ***indicates P-value < 0.001; ****indicates P-value < 0.0001. Scale bars: 50 μm.

## Discussion

Our previous work revealed that Mycn regulates intestinal development in zebrafish through mTOR signalling-mediated ribosomal biogenesis, and activation of mTOR signalling can fully rescue the intestinal deficiency in *mycn* mutants^23^. Our current work emphasized the regulatory function of Mycn on vascular development, especially angiogenesis of the trunk and brain region. We demonstrated that the PI3K signalling regulates the vascular development downstream of Mycn in zebrafish, as activation of PI3K signalling can rescue the vascular development deficiency in *mycn* mutants.

Previous studies have reported that increased expression of the Mycn oncogene in human neuroblastoma cells leads to the downregulation of genes that inhibit cell proliferation, potentially promoting tumor angiogenesis^10^. Our work utilized zebrafish as research models, and demonstrate the essential role of Mycn on angiogenesis during development, revealing a conserved regulatory role of Mycn on the vascular development across different species. More importantly, Targeting the PI3K signalling may act as a therapeutic approach for neuroblastoma treatment in human. Interestingly, in a mouse model of pediatric neurological cancer, application of PI3K inhibitors reduced Mycn expression and induced apoptosis in Mycn-driven cancer cells^34^. In another study focusing on Mycn-amplified neuroblastoma, treatment with the PI3K/mTOR inhibitor NVP-BEZ235 decreased MYCN protein levels and led to the inhibition of VEGF transcription and secretion, disrupting angiogenesis and impeding tumor proliferation^35^. However, our study demonstrates that activation of PI3K signalling failed to rescue the angiogenesis in the head region. These results imply that Mycn’s regulation of angiogenesis in normal cerebral vascular development and neuroblastoma progression may rely on distinct signalling pathways.

Previous works have reported that vascular endothelial growth factors (Vegfs) are essential to specifically promote the brain vessel formation in the zebrafish^36–37^. We also found the Vegfs genes: *vegfba*, *vegfbb* and *vegfc*, were dramatically reduced in *mycn* mutants (Fig. S6). Thus, elevating VEGF levels might aid in alleviating the cerebral vascular development deficiency observed in the mutants. But, how Mycn and VEGFs co-function on cerebral vascular development need to be investigated in the future work.

The processes of vascular and hematopoietic development in zebrafish are closely interconnected, occurring at closely related times and locations. In the zebrafish embryo, vascular development starts around 12 hpf, when lateral plate mesodermal cells differentiate into hemangioblasts. These hemangioblasts have the potential to develop into both hematopoietic cells and vascular endothelial cells^38–39^. By 30 hpf, hematopoietic stem and progenitor cells (HSPCs) emerge from the hemogenic endothelium, a specific group of endothelial cells located in the ventral wall of the dorsal aorta. This process involves an endothelial-to-hematopoietic transition (EHT)^40^. In our current work, we found some *mycn*^+^*kdrl*^+^ cells emerged in the ventral wall of the dorsal aorta at about 30 hpf. Furthermore, the morphology and migration behaviours of those cells are reminiscent of HSPCs^41^. It would be intriguing for future studies to explore whether Mycn plays a role in the production of HSPCs through the EHT.

## Materials and methods

### Animal ethics

Zebrafish strains were conducted following standard procedures, and experimental procedures were approved by The Animal Ethics Committee of the School of Medicine, Zhejiang University. The protocol number is NO. 24278.

### Zebrafish Maintenance and Embryo Collection

The AB strain was used as wild-type zebrafish in this study. The *mycn*-mutant line and transgenic zebrafish line Ki(*mycn*:EGFP) have been described previously^23^. The following transgenic lines were used: Tg(*fli1a*:EGFP) and Tg(*kdrl*:mCherry). All mature zebrafish were maintained under a 14-hour light/10-hour dark cycle at 28℃ in an aquarium supplied with flow water and regular aeration. Zebrafish embryos were obtained by natural spawning, and the sexually mature female and male zebrafish were transferred to a spawning tank at a ratio of 1:1 or 1:2 and separated by a transparent divider. The next morning, fertilized embryos were obtained 20 min after the divider was removed. After washing and disinfecting, the fertilized embryos were transferred into the zebrafish embryo culture buffer −0.3× Danieau Buffer and placed in a 28 ℃ light incubator under controlled light and constant temperature conditions. Embryos were staged according to standard criteria hours post fertilization (hpf) or by days post fertilization (dpf).

### Live imaging

To improve imaging of zebrafish embryos, they were anesthetized in 1% tricaine (BBI). To facilitate the visualization of blood vessels, 50× 1-phenyl-2-thiourea (PTU; Sigma-Aldrich) was added to 0.3× Danieau Buffer at a final concentration of 1 × PTU to inhibit the formation of pigmen. The embryos were placed in a 35 mm glass-bottom culture dish (Cellvis, D35-10-0-N) using 0.25% low melting point agarose (BBI). After 5 min of solidification, the embryos were covered with 0.3× Danieau Buffer containing 1% tricaine and 1× PTU. Time-lapse imaging was performed using an OLYMPUS CSU-W1 spinning-disk confocal microscope with 4×/20× objective lens or 60× oil immersion lens. Z-stacks (Z-step 7 μm or 3 μm) were used to detect the maximum projection intensity. Image processing was performed using cellSens software.

### Whole-mount in situ hybridization (WISH)

The coding sequence of *kdrl* was amplified by PCR, and subsequently cloned and inserted into the PBSK vector. The plasmid was digested with EcoRⅠ (Themo), and probes were transcribed using T7 (Promega) with digoxigenin labeling mix (Roche). For zebrafish embryos, after fixation with 4% paraformaldehyde (PFA; BBI) at 4°C for at least 12 h, embryos were permeated with bleaching solution containing 30% H_2_O_2_ and 5% KOH until the pigment disappeared, and gradient elution was performed using methanol (Sinopharm) and PBS. After that, embryos were stored in 100% methanol (Sinopharm) at −20°C overnight and rehydrated using successive dilutions of methanol with PBST, and washed 4 times with PBST for 5 min per wash. Then, embryos were digested with 10 μg/mL proteinase K for 30 min. After brief fixation with 4% PFA and washing with PBST 4 times for 5 min per wash, embryos were prehybridized in hybridization buffer-HM^+^ (ddH_2_O with 50% formamide, 5× SSC, 500 μg/mL tRNA, 50 μg/mL heparin, 1M citric acid pH 6.0 and 0.1% Tween 20) at 68 °C for least 2 h, and then hybridized with DIG-labeled riboprobe overnight at 68 °C in a rotating hybridization oven. Unbound and excess probe was removed by gradient elution of HM^-^ (HM^+^ without heparin and tRNA) and 2×SSC at 68 °C, followed by two washes in 0.2× SSC for 40 min. Then, after gradient elution of 0.2× SSC and PBST, embryos were incubated with incubation mix (2% bovine serum albumin in PBST with 2% sheep serum) for 3 h at room temperature and then blocked in blocking solution (Anti-Digoxigenin-alkaline phosphatase-Fab fragments in incubation mix) (1:10000; Roche) overnight at 4°C with gentle shaking. Unbound and excess antibodies were removed by washing with PBST six times for 20 min per wash by gently rocking. After three rinses with staining buffer (ddH2O with 10% 1M pH 9.5 Tris-Hcl, 5% 1M MgCl2, 2% 5M NaCl and 0.1% Tween 20), embryos were stained in NBT/BCIP (BBI) staining buffer shielded from light until the appropriate signal was obtained. The reaction was stopped by washing in PBST three times for 5 min per wash. The stained embryos were mounted in glycerol (General reagent) for observation and imaging under a Leica DFC7000 T Microscope.

### TUNEL assay and EdU assay

For TUNEL staining, embryos fixed in 4% PFA at 4°C overnight were treated with ice-cold acetone at −20°C for 30min, rinsed four times in PBST for 20 min, and permeabilized with permeabilization solution (0.1% Triton X-100 and 0.1% sodium citrate in ddH_2_O with 1× PBS) for 2 h. Then, an In Situ Cell Death Detection Kit (Vazyme) was used in accordance with the manufacturer’s instructions. For EdU staining, embryos were dechorionated and cultivated in 500 μM EdU working solution from 60 hpf to 72 hpf. Thereafter, embryos were immersed in 4% PFA at 4°C overnight for fixation. Antigen retrieval was performed with 150 mM tris-HCl (pH 9.0) at 70°C for 15 min. Then, embryos were treated with ice-cold acetone at −20°C for 30 min, rinsed three times in PBSTr buffer for 30 min, blocked in blocking solution (10% bovine serum albumin in PBSTr with 2% goat serum) for 2 h, and rinsed four times in PBST buffer for 20 min. Then, the BeyoClick™ EdU Cell Proliferation Kit with Alexa Fluor 555 (Beyotime) was used in accordance with the manufacturer’s instructions. The specific fluorescence was captured and imaged using confocal laser scanning microscopy (OLYMPUS CSU-W1) with a 20× objective lens. The images were acquired as confocal z-stacks with 7-μm z-steps and projected using maximum intensity.

### Chemical treatment

L-Leu was used to elevate the mTOR pathway. Embryos were injected with 20 mM L-Leu (MCE) into the yolk at 30 hpf, and then imaged at 72 hpf. 740Y-P was used to activate the PI3K pathway. Embryos were exposed to 1mM 740Y-P(MCE) in 0.3× Danieau buffer containing 1× PTU from 10 hpf to 72 hpf.

### Statistical analysis

The area of SIV were quantified in the yolk region of zebrafish embryos across 6-7 somites, and the SIV formation area was measured using ImageJ software. The number of TUNEL-positive cells and EdU-positive cells were counted using ImageJ software. Statistical significance was calculated using independent sample t-tests in GraphPad Prism version 9.5.1. The data are presented as mean ± standard deviation from representative experiments. The statistical parameters, including sample sizes, number of replicate experiments or other quantifications, are indicated for each experiment and shown in figures or figure legends.

### Gene co-expression analysis using the single-cell RNA-seq dataset

The published single-cell RNA-seq dataset of wild-type zebrafish embryos across multiple developmental stages^24^ was employed to investigate the co-expression of *mycn* and *kdrl* (or *fli1a*) in the vasculature system. The vasculature cells were first subsetted from the dataset. Then for each time point of interest, UMAP was used to visualize the vasculature cells, with the *mycn*-positive, *kdrl*-positive (or *fli1a*-positive) and *mycn*&*kdrl*-positive (or *mycn*&*fli1a*-positive) cells labeled in distinct colours.

### Analysis of differential abundance in cell types

The single-cell RNA-seq datasets of 72-h wild-type and *mycn*-mutant embryos were from our previous work^23^. Milo^42^ was employed to analyze the differential abundance in cell types between wild-type and *mycn*-mutant embryos. In brief, a k-nearest neighbour (KNN) graph was constructed and cells were assigned to the defined representative neighbourhoods. The cells belonging to wild-type and *mycn*-mutant embryos in each neighbourhood were counted. Analysis of differential abundance in neighbourhoods was performed using the negative Binomial generalized linear model (GLM) with p-values corrected by the Spatial FDR correction. Each neighbourhood then was assigned a cell type label by finding the most abundant cell type within cells in each neighbourhood. A beeswarm plot was used to visualize the distribution of differential abundance fold changes in different cell types.

### Functional analysis

In the 72-hpf integrated scRNA-seq datasets, the blood vessel cells in *mycn^-/-^* embryo were compared to the wild-type blood vessel cells, and the differentially-expressed (DE) genes were identified through Seurat (https://github.com/satijalab/seurat) with a Bonferroni adjusted p-value < 0.05. Down regulated genes in the *mycn^-/-^* blood vessel cells were imported into Metascape (https://metascape.org) for functional analysis.

## Additional information

Supplemental Information: Supplementary Figures 1-6 and Supplementary Movies 1-5.

## Acknowledgments

We would like to thank the members of Laboratory of Development and Organogenesis (LDO) at Zhejiang University for helpful suggestions and discussions. We thank Shuang-Shuang Liu from the Imaging Platform and Ying-Niang Li from the zebrafish core facility at Zhejiang University School of Medicine for their technical support. This work was supported by grants from the Chinese National Key Research and Development Project (2022YFA1103100, 2019YFA0802402) and the National Natural Scientific Foundation of China (32050109), in part by the Guizhou Province’s Science and Technology Major Project (QianCXTD[2021]002, GCC[2023]034).

## Author Contributions

PFX, LPS and YF conceived and designed the research; GQZ, TC, PYW, JM, FY, YD and YFL performed research; TC and YD analyzed the single-cell RNA-seq and bulk RNA-seq data; GQZ, TC and PFX wrote the manuscript; all authors reviewed and approved the manuscript. LPS, PFX and YF supervised this study.

